# Low-pass sequencing plus imputation using avidity sequencing displays comparable imputation accuracy to sequencing by synthesis while reducing duplicates

**DOI:** 10.1101/2022.12.07.519512

**Authors:** Jeremiah H. Li, Karrah Findley, Joseph K. Pickrell, Kelly Blease, Junhua Zhao, Semyon Kruglyak

## Abstract

Low-pass sequencing with genotype imputation has been adopted as a cost-effective method for genotyping. The most widely used method of short-read sequencing uses sequencing by synthesis (SBS). Here we perform a study of a novel sequencing technology — avidity sequencing. In this short note, we compare the performance of imputation from low-pass libraries sequenced on an Element AVITI system (which utilizes avidity sequencing) to those sequenced on an Illumina NovaSeq 6000 (which utilizes SBS) with an SP flow cell for the same set of biological samples across a range of genetic ancestries. We observed dramatically lower duplication rates in the data deriving from the AVITI system compared to the NovaSeq 6000, resulting in higher effective coverage given a fixed number of sequenced bases, and comparable imputation accuracy performance between sequencing chemistries across ancestries. This study demonstrates that avidity sequencing is a viable alternative to the standard SBS chemistries for applications involving low-pass sequencing plus imputation.

## Introduction

Low-pass whole-genome sequencing followed by imputation has become increasingly popular as a cost-effective genotyping method for large-scale genetic studies in both humans and other species (Li et al., 2021; Wasik et al., 2021; Tran et al., 2020; Rubinacci et al., 2021; Nosková et al., 2021; Li et al., 2022; Buckley et al., 2022; Martin et al., 2021).

In this study, we wished to evaluate the utility of short-read sequencing data generated using a recently introduced method termed avidity sequencing for the application of imputation to large haplotype reference panels.

Avidity sequencing is described in Arslan et al. (2022). Briefly, linear library molecules are circularized through hybridization to splint oligos followed by ligation. The circularized library molecules are captured on a flow cell and amplified via rolling circle amplification, forming concatemer template strands. Sequencing cycles are initiated following hybridization of sequencing primers.

The sequencing cycles differ from sequencing by synthesis (SBS) in several important ways. First, the process of stepping along the DNA template strand is separated from the process of nucleotide identification. This decoupling enables the independent optimization of each step with respect to quality and reagent consumption. Second, a dye-labeled polymer with multiple, identical nucleotides attached, termed an avidite, is used for base identification. In the presence of a polymerase evolved for this purpose, the avidite is able to bind multiple DNA copies within the concatemer. The multivalent binding is highly specific and enables a reduction in the concentration of reporting nucleotides from micromolar to nanomoloar due to a negligible dissociation rate. Four avidites, each corresponding to a specific nucleotide and dye label, are used to identify nucleotides in the DNA template.

In previous work, we compared the performance of imputation from low-pass whole-genome sequencing (lpWGS) from Illumina and BGI machines to that from a commonly used genotyping array, the Illumina GSA, in the context of statistical power for genome-wide association studies and the estimation of polygenic risk scores (Li et al., 2021).

We performed a similar study here to evaluate the performance of imputation from low-pass sequence data for a set of diverse individuals sequenced on both the Element AVITI system and the Illumina NovaSeq 6000 using an SP flow cell to the same target coverage of 0.8×.

Previously, we found that a key predictive metric of imputation performance (as measured by non-reference concordance to a held-out gold standard) for sequencing data was the *effective coverage* of a sample, which is defined as a function of the fraction of polymorphic sites in a haplotype reference panel covered by at least one sequencing read (Methods) (Li et al., 2021). This metric is useful because it gives an indication not only of the overall nominal sequencing coverage (*i.e*., the number of bases sequenced divided by the genome size of the organism), but also takes into account the evenness of genome-wide sequencing coverage.

In this study, we observed overall lower duplication rates in the data deriving from the AVITI system compared to the NovaSeq 6000, resulting in higher effective coverage given a fixed number of sequenced bases, and comparable imputation accuracy performance between sequencing chemistries across ancestries, suggesting that avidity sequencing provides a viable alternative to SBS for applications involving lpWGS followed by imputation.

## Results

### Experimental Design

In order to compare the relative performance of low-pass sequencing across populations, we selected 48 individuals of European ancestry and 48 individuals of African ancestry from the 1000 Genomes Phase 3 release (1KGP3) (1000 Genomes Project Consortium, 2015) for sequencing at a target 0.8× coverage on the NovaSeq 6000 (with an SP flow cell) and the AVITI system.

In order to make the experiment as controlled as possible, we prepared a single library pool from the same set of gDNA samples before dividing it for parallel sequencing on both machines (Methods).

For each sample successfully sequenced and passing QC, we imputed the sequence data to the 1KGP3 haplotype reference panel in a leave-one-out manner (Methods) and compared the imputed calls to the left-out gold-standard genotypes from the reference panel. Specifically, the following comparisons were performed on bi-allelic autosomal SNPs.

### Data Quality Control

Of the 96 samples sequenced via each chemistry, 93 passed QC in both datasets. All analyses below are restricted to this subset of individuals.

Overall, we observed comparable nominal coverages, with a mean (standard deviation) of 1.09× (0.36) for the AVITI data and 0.95× (0.29) for the NovaSeq data.

Notably, however, the duplication rate for the libraries differed substantially across the two sequencing chemistries, with the AVITI data having a twenty-fold lower average duplication rate compared to the NovaSeq data. Specifically, we observed a mean (standard deviation) duplication rate of 0.27% (0.058%) for the AVITI and 5.5% (0.57%) for the NovaSeq (Fig. 1a). A two-sample unpaired Welch’s *t*-test for differences in means in duplication rate between the AVITI and NovaSeq data yielded statistically significant (*P* <0.001) results.

**Figure 1:**
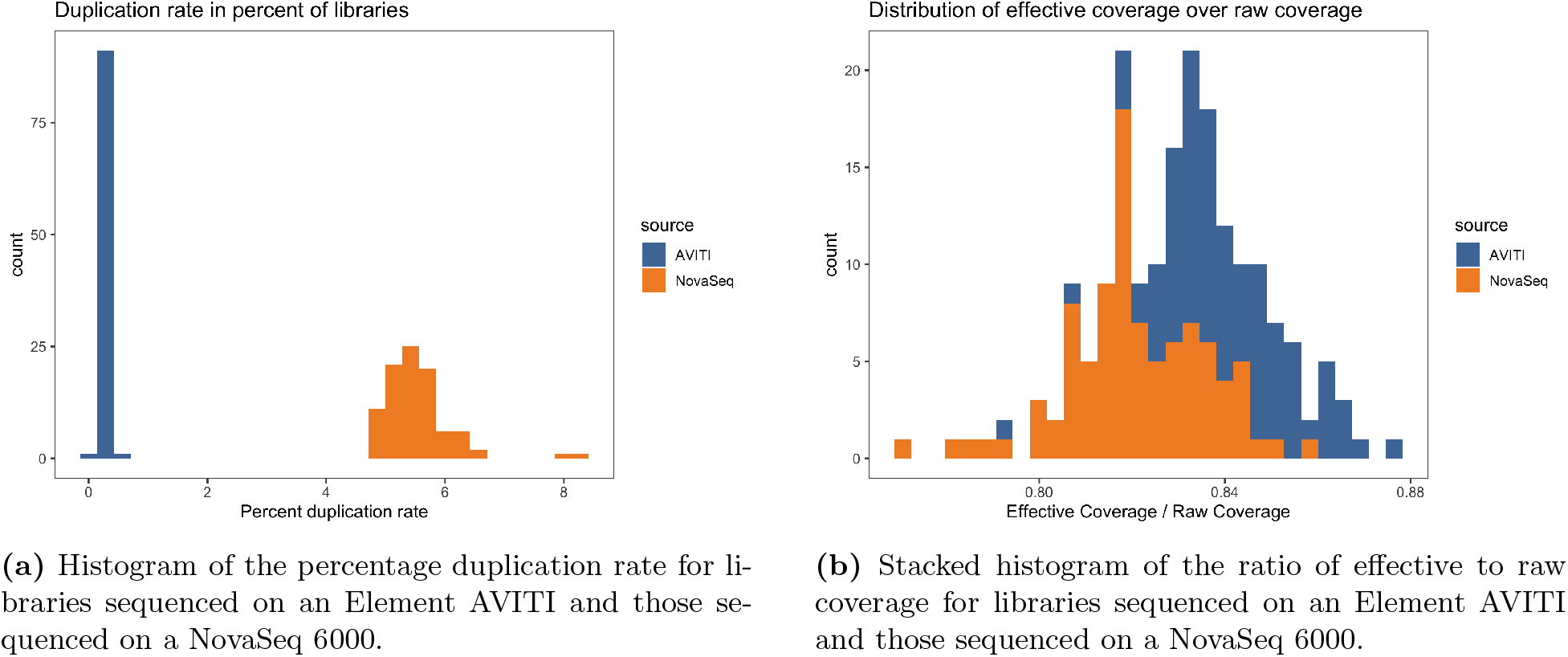
Distribution of measures of duplication rate (a) and sequencing evenness (b).

Since duplicate reads are redundant, a higher rate of duplication should correspond to less spatially uniform coverage (with respect to the genomic coordinate system) of the whole genome (compared to Poisson sampling) given a fixed number of unfiltered sequence reads. As a result, for a given nominal coverage (that is, the number of sequenced bases divided by the genome size), the sequencing chemistry with a lower duplication rate should yield a higher *effective coverage* (denoted λ_eff_) (Methods), *ceteris paribus*, since λ_eff_ is a function of the proportion of sites in a reference panel covered by at least one read (that is, λ_eff_ is unaffected by the number of reads at a site above one).

As Figure 1b demonstrates, this is indeed the case, as the distribution of the ratio of effective to raw coverage (higher is better) is shifted right for the AVITI data compared to the NovaSeq data (with statistically significantly different means of 0.84 and 0.82 respectively; Welch’s *t*-test *P* <0.001). The effect is not dramatic, however, particularly when considering the fold difference between raw duplication rates between the sequencing chemistries.

We note here that the λ_eff_ is particularly relevant to low-pass applications as the input data for imputation from low-pass sequencing are precisely those reads which overlap sites in the reference panel; for more details, see Li et al. (2021).

### Comparison of imputation quality metrics across sequencers

To evaluate imputation performance, we computed the non-reference concordance (NRC) of each imputed sample’s genotypes to the held-out “truth” data. Consistent with previous results, we observed that imputed genotypes in the EUR cohort were on average more accurate than in the AFR cohort, though the difference is not large. For the AVITI data, the mean (standard deviation) NRC was 0.977 (0.00396) in the EUR cohort and 0.970 (0.00474) in the AFR cohort. For the NovaSeq data, the mean (standard deviation) NRC was 0.976 (0.00429) in the EUR cohort and 0.968 (0.00501) in the AFR cohort.

We also examined NRC across the allele frequency spectrum, and observed that results were comparable across sequencing chemistries. Figure 2 depicts the mean NRC in a given allele frequency bin (where the allele frequencies are drawn from the 1KGP3) for a given superpopulation and sequencing chemistry. Consistent with previous observations (Marchini, 2019; Li et al., 2021), imputation performance at low allele frequencies is higher in the AFR cohort than the EUR cohort, with the pattern reversing at higher allele frequencies. As before, we hypothesize that these represent the extremes of two different allele frequency regimes, wherein at lower allele frequencies, the greater genetic diversity in Africans dominates and affords more accurate rare variant tagging and thus increased imputation accuracy, and at higher allele frequencies, the stronger linkage disequilibrium structure within European populations dominates and thus affords increased imputation accuracy relative to African populations.

**Figure 2:**
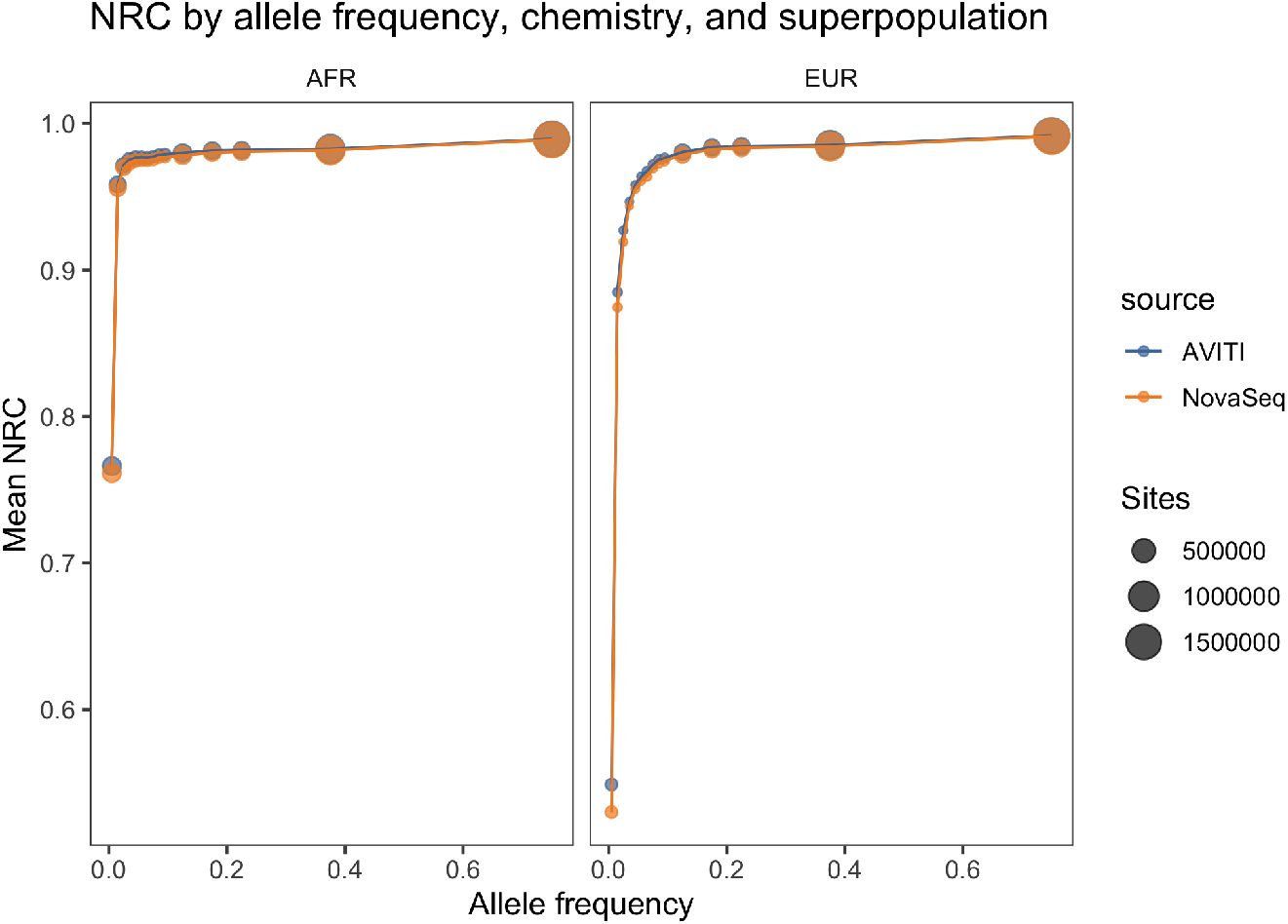
Non-reference concordance (NRC) across the allele frequency spectrum from the 1KGP3, stratified by chemistry and superpopulation. Note that the NRC at lower allele frequencies is higher on average for the AFR cohort and that results are comparable across sequencing chemistries. Note also that the number of sites evaluated is lower at lower allele frequencies due to the fact that NRC is computed only on sites where both the imputed and gold-standard callsets are called homozygous reference.

We therefore see comparable imputation accuracy between the sequencing chemistries even with the large difference in duplication rates. We hypothesize that the effect of lower duplication rates on the accuracy of imputation is more pronounced at lower coverages, where “every read counts” to a greater extent than it does at this coverage regime.

## Discussion

For the last decade, sequencing-by-synthesis has been the predominant method underlying high-throughput short-read whole-genome-sequencing. Recently, a number of alternative methods such as avidity sequencing and sequencing-by-binding have been proposed for highly parallel high-throughput sequencing (Arslan et al., 2022; Pacific Biosciences, 2022). In this study, we examined the viability of avidity sequencing as an alternative to SBS in the context of low-pass whole genome sequencing followed by imputation.

We selected cohorts of EUR and AFR individuals from the 1KGP3 project and prepared libraries using the same preparation method before dividing the library pool for sequencing to the same target coverage of 0.8× on an Element AVITI system and an Illumina NovaSeq 6000 with an SP flow cell.

We observed that the duplication rate within the dataset deriving from avidity sequencing was dramatically (twenty-fold) lower than the dataset deriving from the SBS method, and that the ratio of effective coverage to nominal coverage for the former dataset was higher on average than the latter, though the effect size was considerably smaller. We observed excellent imputation accuracy (as measured by non-reference concordance to a gold standard) across the allele frequency spectrum and comparable results from both datasets, with an average NRC in the EUR cohort of 0.977 for the AVITI and 0.976 for the NovaSeq.

We conclude that avidity sequencing presents a viable alternative to SBS methods for the purposes of low-pass whole genome sequencing followed by imputation.

We hypothesize that lower duplication rates may be most relevant for *ultra-low* (*e.g*., <0.1×) coverage applications, where “every read counts.”

For instance, in large-scale agricultural applications where hundreds of thousands of individuals are genotyped every year and where sequencing costs represent a nontrivial proportion of the total cost of a genomic prediction program, reducing the proportion of redundant reads (*i.e*., duplicates) would allow for even higher levels of multiplexing during a single sequencing run, resulting in a lower marginal genotyping cost.

## Methods

### Data Generation

Purified genomic DNA (gDNA) from 48 selected individuals of European ancestry and 48 selected individuals of African ancestry from the 1000 Genomes Project Phase 3 was obtained from the NIGMS Human Genetic Cell Repository at the Coriell Institute for Medical Research. Genomic DNA was extracted from immortalized B lymphocytes and eluted in TE buffer (10 mM Tris, pH 8.0/1 mM EDTA) for shipping.

To prepare the 96 gDNA samples for sequencing, 30 *μ*L from each was plated and diluted to 10 ng/*μ*L using 10 mM Tris-HCl, and sequencing libraries were prepared using a miniaturized version of the KAPA HyperPlus kit (Roche KK8514) with Illumina-compatible KAPA Unique Dual-Indexed Adapters (Roche KK8727).

Following library preparation, two multiplexed library pools were prepared for parallel sequencing on an Illumina instrument and an Element instrument. To multiplex the libraries for sequencing, a subset volume of 6 *μ*L was pooled from all libraries to prepare two identical pools containing 96 samples each. The pooled libraries were purified with a double-sided size-selection using SPRIselect paramagnetic beads (Beckman Coulter Life Sciences B23318) to narrow the library fragment size range and remove dimerized adapters from the samples. To characterize the purified pools, the concentration was measured using the Invitrogen Qubit Fluorometer (Thermo Fisher Scientific Q33238), and the library fragment size was assessed using the Agilent 2100 Bioanalyzer (Agilent G2939BA) with a High Sensitivity DNA Kit (Agilent 5067-4626).

Sequencing was performed in parallel to 0.8× target coverage (∼8-9M reads at 2×150bp) for each sample using both the Illumina NovaSeq 6000 and the Element AVITI System. The Illumina sequencing run was performed using the SP flow cell and paired-end 150bp reads. For the samples to be run on the Element AVITI, 0.5 pmol (30 *μ*L of 16.7nM) of the pooled libraries were processed in a single reaction using the Adept Compatibility Workflow Kit (Element Biosciences, Cat#830-00003). The final circularized library was quantified using qPCR standard and primer mix provided in the Adept Compatibility Workflow Kit. The quantified library was denatured and sequenced on Element AVITI system using 2×150 paired end reads.

### Effective Coverage

The *effective coverage* λ_eff_ of a sample having undergone low-pass sequencing plus imputation to a haplotype reference panel is defined as

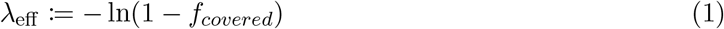

where *f_covered_* is the fraction of sites in the reference panel covered by at least one sequencing read. For a more detailed derivation and background on this quantity, see Li et al. (2021).

### Imputation

Each pair of FASTQs was aligned to the `hs37-1kg` reference assembly (obtained from NCBI at the following URL: ftp://ftp-trace.ncbi.nih.gov/1000genomes/ftp/technical/reference/human_g1k_v37.fasta.gz). From the resulting alignments, we extracted pileups using `samtools mpileup` and computed genotype likelihoods at all sites in the 1KGP3 haplotype reference panel using a custom program (1000 Genomes Project Consortium, 2015; Danecek et al., 2021). Thereafter, these genotype likelihoods were fed into a custom hardware-accelerated version of `GLIMPSE` and genotypes were independently imputed for each sample to the 1KGP3 haplotype reference panel in a leave-one-out manner (Rubinacci et al., 2021). Variants for which none of the posterior probabilities for the three possible genotypes (homozygous reference, heterozygous, homozygous alternative) was greater than 0.9 were marked as low confidence and filtered out before the comparisons performed above.

### Non-Reference Concordance

The *non-reference concordance* at the intersection of sites between two genotype callsets is defined as the ratio of the number of sites with matching genotypes (less the sites at which both callsets have a homozygous reference call) over the total number of sites in the intersection (less the sites at which both callsets have a homozygous reference call). That is, it is the genotype concordance calculated at all sites which do not have a homozygous reference call for both callsets. This metric is preferred over “overall concordance” given that for a given individual, the vast majority of genotype calls at sites within a large haplotype reference panel will be homozygous reference.

## Data access

The raw sequence data generated in this study have been submitted to the NCBI BioProject database (https://www.ncbi.nlm.nih.gov/bioproject/) under accession number PRJNA909799. All raw sequence data, alignments, and imputed variants are also available in a public AWS S3 bucket at

~~~
s3://gencove-element-paper-2022.
~~~

## Acknowledgements

J.H.L., J.K.P, and S.K. conceived of the study. J.H.L conducted the analyses. K.F., K.B., J.Z. were responsible for the experimental work and data acquisition.

We thank Chase Mazur for contributions to the manuscript regarding the preparation of samples.

We thank Lex Flagel, Gillian Belbin, and Yuwan Lam for comments on previous drafts of this manuscript.

